# Molecular epidemiology of *Escherichia coli* producing CTX-M and plasmid AmpC-type β-lactamases from dairy farms identifies a dominant plasmid encoding CTX-M-32 but no evidence for transmission to humans in the same geographical region

**DOI:** 10.1101/845917

**Authors:** Jacqueline Findlay, Oliver Mounsey, Winnie W.Y. Lee, Nerissa Newbold, Katy Morley, Hannah Schubert, Virginia C. Gould, Tristan A. Cogan, Kristen K. Reyher, Matthew B. Avison

**Affiliations:** School of Cellular & Molecular Medicine, Biomedical Sciences Building, University of Bristol, University Walk, Bristol, UK; Bristol Veterinary School, University of Bristol, Langford, UK

## Abstract

Third-generation cephalosporin resistance (3GC-R) in *Escherichia coli* is a rising problem in human and farmed animal populations. We conducted whole genome sequencing analysis of 138 representative 3GC-R isolates previously collected from dairy farms in South West England and confirmed by PCR to carry acquired 3GC-R genes. This analysis identified *bla*_CTX-M_ (131 isolates: encoding CTX-M-1, −14, −15, −32 and the novel variant, CTX-M-214), *bla*_CMY-2_ (6 isolates) and *bla*_DHA-1_ (one isolate). A highly conserved plasmid was identified in 73 isolates, representing 27 *E. coli* sequence types. This novel ~220 kb IncHI2 plasmid carrying *bla*_CTX-M-32_ was sequenced to closure and designated pMOO-32. It was found experimentally to be stable in cattle and human transconjugant *E. coli* even in the absence of selective pressure and was found by multiplex PCR to be present on 26 study farms representing a remarkable range of transmission over 1500 square kilometres. However, the plasmid was not found amongst human urinary *E. coli* we have recently characterised from people living in the same geographical location, collected in parallel with farm sampling. There were close relatives of two *bla*_CTX-M_ plasmids circulating amongst eight human and two cattle isolates, and a closely related *bla*_CMY-2_ plasmid found in one cattle and one human isolate. However, phylogenetic evidence of recent sharing of 3GC-R strains between farms and humans in the same region was not found.

**Importance:** Third-generation cephalosporins (3GCs) are critically important antibacterials and 3GC-resistance (3GC-R) threatens human health, particularly in the context of opportunistic pathogens such as *Escherichia coli*. There is some evidence for zoonotic transmission of 3GC-R *E. coli* through food, but little work has been done examining possible transmission (e.g. via interaction of people with the local near-farm environment). We characterised acquired 3GC-R *E. coli* found on dairy farms in a geographically restricted region of the United Kingdom and compared these with *E. coli* from people living in the same region, collected in parallel. Whilst there is strong evidence for recent farm-to-farm transmission of 3GC-R strains and plasmids – including one epidemic plasmid that has a remarkable capacity to transmit – there was no evidence that 3GC-R found on study farms had a significant impact on circulating 3GC-R *E. coli* strains or plasmids in the local human population.

## Introduction

Third-generation cephalosporin-resistant (3GC-R) *Escherichia coli* have been increasingly reported in both animal and human populations and are considered pathogens of major concern for humans (1,2). 3GCs such as cefotaxime and ceftazidime have been listed by the World Health Organisation (WHO) as “highest-priority critically important antimicrobials” (HP-CIAs) because of their importance for human health (3). Resistance to 3GCs in *E. coli* can be caused by several mechanisms but is primarily attributed to the acquisition of Extended Spectrum β-Lactamases (ESBLs) or plasmid-mediated AmpC β-lactamases (pAmpCs) (4). Plasmids encoding ESBLs and pAmpCs frequently harbour additional resistance genes and so can present a significant therapeutic challenge (5). In recent years, the promotion and implementation of the ‘One Health’ approach in antimicrobial resistance by the WHO has emphasised the importance of surveillance in both animal and human populations and has highlighted gaps in this knowledge (6). In humans it has been well established in numerous global studies that certain *E. coli* lineages (e.g. *bla*_CTX-M_-encoding ST131) play a major role in the dissemination of ESBL genes, however such a depth of information does not exist for isolates from animal populations (2). Human-associated pandemic lineages have been reported in animal populations albeit to a much lesser extent than in human populations (7).

In humans, *bla*_CTX-M_ variants are the globally dominant ESBL type with some variants exhibiting geographical associations (e.g. *bla*_CTX-M-15_ in Europe and North America and *bla*_CTX-M-14_ in Asia) (2). Transmission of ESBLs occurs largely through horizontal gene transfer, with conjugative IncF plasmids being reported as the dominant vehicles for *bla*_CTX-M_ genes (8,9). Previous studies using typing methodologies including WGS have suggested transmission of both strains and ESBL plasmids across animal and human populations (10,11). Epidemic plasmids have been reported across different host populations and in multiple countries (12). For example, one epidemic plasmid type – pCT, encoding *bla*_CTX-M-14_ – was identified in cattle and human *E. coli* isolates in England and found to exist in human isolates from several countries across 3 continents (12).

Antimicrobial use in food animals may provide selective pressure for resistance genes/plasmids which could theoretically be spread to humans (13). However, recent reports suggest that such transmission is limited, at least in the UK (14). In dairy farming, antibiotics are used both therapeutically in the treatment of common infections such as mastitis, and preventatively. For example, in so-called dry cow therapy, an antibacterial preparation inserted into a cow’s udder between lactations to prevent mastitis (15). A survey of dairy farms in England and Wales in 2012 revealed that the fourth-generation cephalosporin (4GC) cefquinome (another HP-CIA) was the most used dry cow therapy treatment (16). By 2017, however, only 5.3% of total dry cow therapy active ingredients were HP-CIAs. Indeed, there has been a significant decline in the use of HP-CIAs on dairy farms in the UK in recent years (17).

Recently, we reported a survey of 53 dairy farms located in South West England where we investigated the prevalence of 3GC-R *E. coli* (18). From 1226 such isolates, PCR analysis confirmed that 648/1226 (52.7%) carried *bla*_CTX-M_ and 13/1226 (1.1%) carried a pAmpC gene. The remaining 566/1226 (46.2%) isolates did not carry any putative acquired 3GC-R gene, and a subsequent analysis confirmed that hyper-production of the chromosomally encoded AmpC β-lactamase was the mechanism of 3GC-R in these isolates (19).

The study reported here aimed to characterise, using WGS, representative *bla*_CTX-M_ and pAmpC-positive *E. coli* isolates collected on dairy farms during our earlier surveillance study (18). Furthermore, our aim was to compare these isolates at strain and plasmid-encoded 3GC-R gene level with *bla*_CTX-M_ and pAmpC positive urinary *E. coli* isolates (20) collected from humans living in the same 50 x 50 km region that was the location of the majority of dairy farms under study.

## Results and Discussion

### WGS analysis of *E. coli* carrying acquired 3GC-R genes from dairy farms

One hundred and thirty-eight representative isolates, PCR-positive for *bla*_CTX-M_ or pAmpC genes (18) and chosen to give coverage of all 42 farms positive for acquired 3GC-R genes, were subjected to WGS (**Table 1**). *bla*_CTX-M-32_, encoding a group 1 enzyme first described in a human clinical isolate in 2004 (21), was the most common 3GC-R gene identified and was found in 79/138 sequenced isolates encompassing 27 *E. coli* sequence types (STs) from 25 farms Other 3GC-R genes identified were: *bla*_CTX-M-14_ (18 isolates, 6 STs from 9 farms), *bla*_CTX-M-1_ (16 isolates, 8 STs from 6 farms), *bla*_CTX-M-15_ (16 isolates, 5 STs from 10 farms), *bla*_CMY-2_ (6 isolates, 3 STs from 3 farms), *bla*_CTX-M-214_ (3 isolates, 2 STs from 3 farms) plus one isolate harbouring *bla*_DHA-1_ and one isolate having both *bla*_CTX-M-1_ and *bla*_CTX-M-14_. CTX-M-214 (GenBank Accession No. MH121688) is a novel CTX-M-9 variant, first identified in this study, which differs from CTX-M-9 by a single amino acid, Ala112Thr. In all three isolates encoding *bla*_CTX-M-214_, the gene was identified on a contig which also encoded an IncI-ST26 plasmid replicon as well as *aadA2*, *sul1*, and *dfrA16*.

**Table 1.**
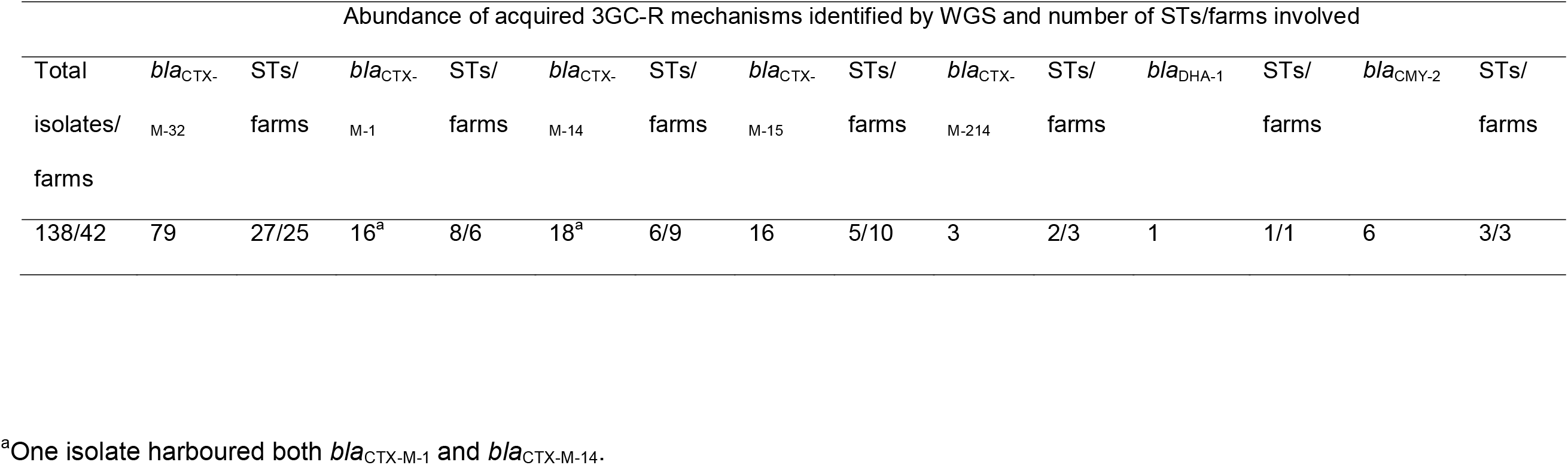
Characteristics of 3GC-R 138 isolates subjected WGS.

### Identification and characterisation of pMOO-32

Following observations of the high prevalence of *bla*_CTX-M-32_, a search for common plasmid replicon types was conducted which revealed an IncHI2-ST2 replicon in almost all the sequenced *bla*_CTX-M-32_-positive isolates. Transconjugations were attempted into *E. coli* DH5α using *bla*_CTX-M-32_-positive farm isolate DK as donor (**Table 2**). One successful transconjugant was sent for WGS employing both long and short read methodologies to sequence the plasmid to closure. pMOO-32 is a 226,022-bp conjugative plasmid belonging to the ST2-IncHI2 incompatibility group, harbouring *repHI2* and *repHI2A* replication genes. It contains 245 putative ORFs and has a GC content of 45.5% (**Figure 1**). pMOO-32 encodes the following antimicrobial resistance genes: *bla*_CTX-M-32_, *strA*, *strB*, *aph(6)-Ic*, *aph(3’)-IIa* and *tetB* as well as genes encoding resistance to the heavy metal compound, tellurite (*terABCDEFWXYZ*) and a HipAB type II toxin-antitoxin system along with a second partial system (*higB* toxin gene). *bla*_CTX-M-32_ is encoded downstream of an IS*ECp1* element within which there is an IS*Kpn26* insertion encoded in the opposite orientation (**Figure 2**). This same genetic environment was also observed in 4 *bla*_CTX-_ M-32-positive but IncHI2 plasmid-negative ST10 isolates collected from 2 farms. There were 2 additional IncHI2 plasmid-negative ST765 isolates, both from the same farm, that carried *bla*_CTX-M-32_ where the immediate genetic environment differed by a truncation in IS*Ecp1*.

**Table 2.** Disc susceptibility testing of *E. coli* DK and pMOO-32 transconjugants of various *E. coli* recipient strains.

| Antimicrobial Agent | Zone Diameters (mm) |  |  |  |  |  |  |  |  |  |  |  |  |  |
| --- | --- | --- | --- | --- | --- | --- | --- | --- | --- | --- | --- | --- | --- | --- |
|  | *E. coli* DK | S/I/R | *E. coli* DH5α | S/I/R | *E. coli* DH5α TR | S/I/R | *E. coli* HG | S/I/R | *E. coli* HG TR | S/I/R | *E. coli* HC4 | S/I/R | *E. coli* HC4 TR | S/I/R |
| AMP | <6 | R | 30 | S | <6 | R | <6 | R | <6 | R | 20 | S | <6 | R |
| CTX | 10 | R | 45 | S | 18 | I | 35 | S | 10 | R | 34 | S | 13 | R |
| CAZ | 20 | I | 45 | S | 30 | S | 34 | S | 20 | I | 32 | S | 23 | S |
| FEP | 19 | R | 45 | S | 28 | S | 28 | S | 18 | R | 35 | S | 24 | I |
| ETP | 30 | S | 45 | S | 44 | S | 34 | S | 33 | S | 36 | S | 36 | S |
| ATM | 15 | R | 45 | S | 24 | I | 34 | S | 15 | R | 35 | S | 19 | R |
| STR<sup>a</sup> | <6 | R | 25 | S | 10 | R | <6 | R | <6 | R | 13 | R | <6 | R |
| TOB | 17 | S | 28 | S | 26 | S | <6 | R | <6 | R | 18 | S | 19 | S |
| NEO<sup>a</sup> | 8 | R | 20 | S | 12 | R | <6 | R | <6 | R | 14 | I | 8 | R |
| TET<sup>a</sup> | <6 | R | 36 | S | <6 | R | <6 | R | <6 | R | 30 | S | <6 | R |

AMP, ampicillin; CTX, cefotaxime; CAZ, ceftazidime; FEP, cefepime; ETP, ertapenem; ATM, aztreonam; STR, streptomycin; TOB, tobramycin; NEO, neomycin; TET, tetracycline; TR, pMOO-32 transconjugant.
<sup>a</sup>Streptomycin and neomycin sensitivities were determined using tobramycin EUCAST interpretation guidelines, and tetracycline according to guidelines for *Yersinia enterocolitica*.

**Figure 1.**
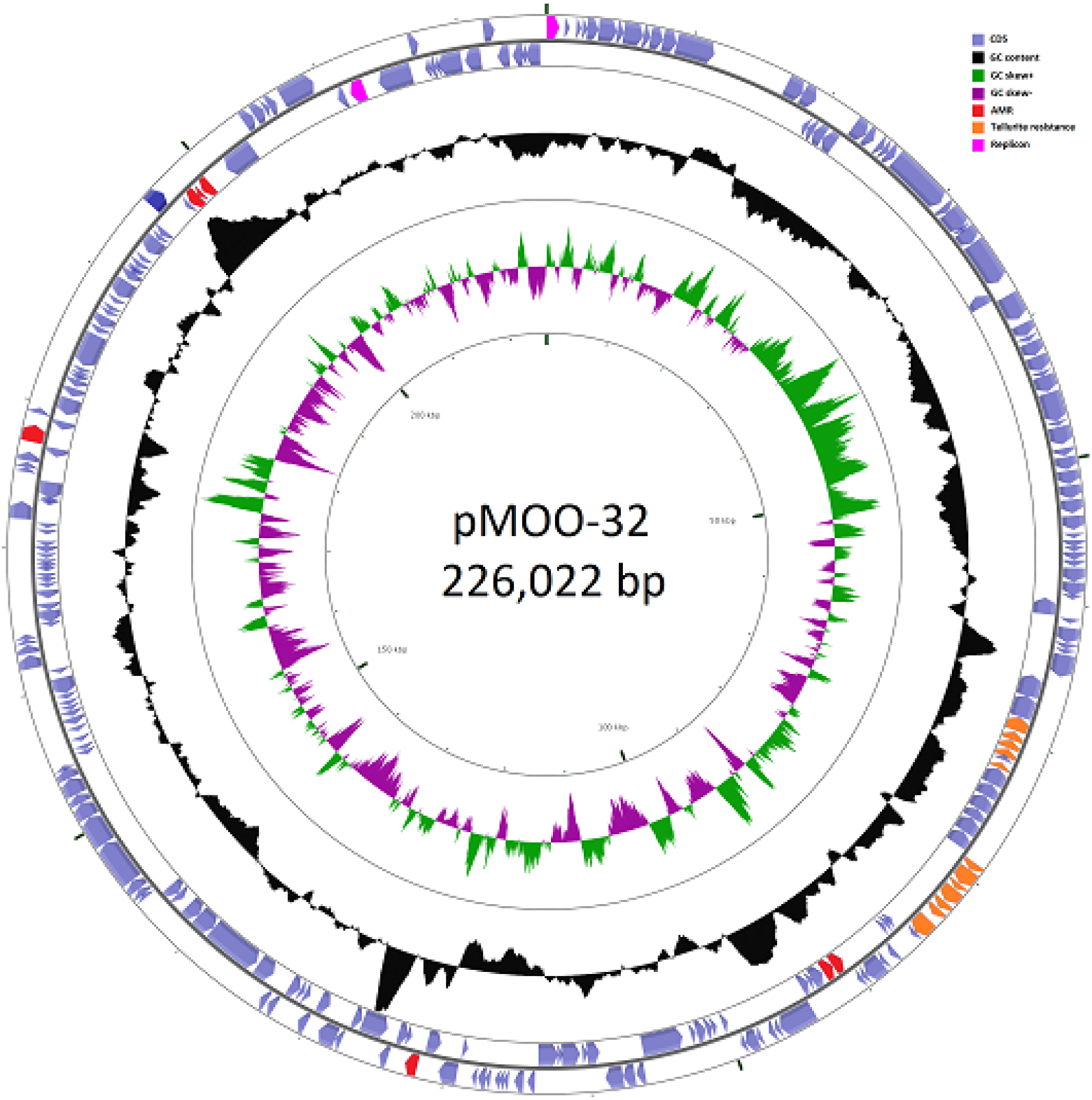
Plasmid pMOO-32 created using CGView (40).

**Figure 2.**
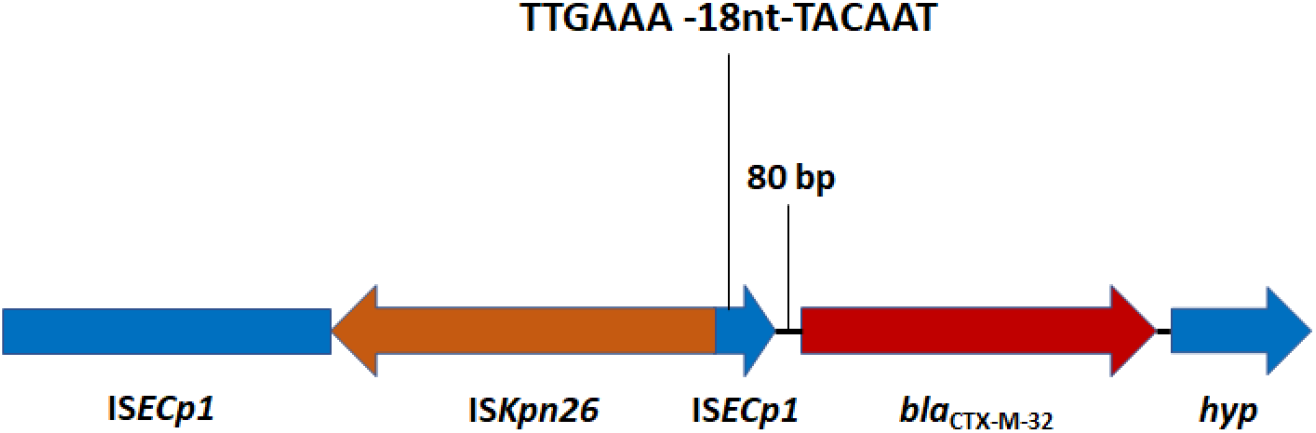
The genetic environment of *bla*_CTX-M-32_ in pMOO-32 and other IncHI2 positive, *bla*_CTX-M-32_-harbouring isolates.

Transconjugation attempts using the pMOO-32-positive farm isolate DK as donor into a 3GC-susceptible (3GC-S) cattle ST88 *E. coli* isolated from one of the study farms (18) as well as into a 3GC-S human urinary ST1193 *E. coli* isolate (20) were both successful (**Table 2**). ST1193 was selected as a recently described fluoroquinolone-resistant global clone, often implicated as a cause of human infections (22), whilst ST88 was selected as a particularly prevalent ST among cattle isolates (23). Antimicrobial disc testing showed that the pMOO-32-carrying donor was, as expected from the genotype, resistant to ampicillin, cefotaxime, cefepime, aztreonam, streptomycin, neomycin, and tetracycline. The cattle ST88 and human ST1193 transconjugants were, additional to their starting wild-type resistance profile, resistant to cefotaxime, cefepime and aztreonam. These results (**Table 2**) are indicative of the functionality of the *bla*_CTX-M-32_ gene harboured by pMOO-32.

### Epidemiology of pMOO-32-like plasmids on farms

The complete nucleotide sequence of pMOO-32 was submitted to GenBank under accession number MK169211. Using this sequence as a reference, sequencing reads from all 73 isolates positive for *bla*_CTX-M-32_ and the IncHI2 replicon were mapped; this indicated that the plasmids exhibited 94-100% identity to the reference sequence. Any differences could be attributed to a loss or gain of mobile genetic elements, but no major rearrangements were observed to the plasmid backbone or changes to resistance gene content. These 73 isolates carrying pMOO-32-like plasmids comprised 27 *E. coli* STs. On 10 farms, pMOO-32-like plasmids were found in isolates of more than one ST. The most frequently identified STs were ST69 and ST10, found in 18 isolates from 7 farms, and 6 isolates from 4 farms, respectively. Therefore, we conclude that pMOO-32-like plasmids are dominant in this study area, largely a result of horizontal rather than clonal transmission.

We subsequently designed a multiplex PCR, based on the pMOO-32 sequence, and used to it screen all group 1 *bla*_CTX-M_-PCR-positive isolates from our study farms (18) not previously subjected to WGS for the presence of pMOO-32-like plasmids. It was found that 26/53 (49.1%) farms within our study area tested positive for the presence of a pMOO-32-like plasmid using this test. The origins and geographical reach of pMOO-32 remain to be established, but all positive farms were located in a 40 × 40 km sub-region of the wider study area, where 26/33 farms were positive, suggesting that the plasmid remained geographically confined at the time of sample collection.

We hypothesised that the observed dominance of pMOO-32-like plasmids could be a consequence of the HipAB-type II toxin-antitoxin system found on the plasmid, leading to persistence even in the absence of antibacterial selective pressure. Growth curve assays indicated a 12-40% fitness cost (reduction in OD_600_) of pMOO-32 carriage in *E. coli* DH5α at the end of the exponential growth phase in M9 minimal medium (data not shown). However, despite this cost in growth terms, pMOO-32 was stably maintained over 10 days of passaging in the absence of antibiotic pressure in the farm isolates, their respective DH5α transconjugants, and the human and cattle *E. coli* isolate transconjugants tested.

### No evidence for recent sharing of 3GC-R *E. coli* and limited evidence for recent sharing of 3GC-R *E. coli* plasmids between humans and dairy farms

The ability of pMOO-32 to readily transfer into human urinary *E. coli* ST1193 (**Table 2**) and be maintained in the absence of antimicrobial pressure indicates the zoonotic potential of this plasmid. We next aimed to see if there was evidence of recent sharing of 3GC-R isolates or plasmids, including pMOO-32, between dairy farms and humans living in the same geographical region as the farms. To do this, we compared WGS data from farm isolates described above with data from human urinary *E. coli* collected in parallel within the same 50 × 50 km geographical region (20). Since 10 dairy farms under study here lie outside the 50 × 50 km region (18), isolates from these farms were excluded.

Core genome phylogenetic analysis of 324 sequenced (112 dairy farm and 212 human) *E. coli* isolates carrying acquired 3GC-R genes collected within the target region (**Figure 3**) revealed only four STs including examples of both farm and human isolates. In no case was the single nucleotide polymorphism (SNP) difference between any pair of human and farm isolate core genomes suggestive of recent transmission. ST10 was the closest (≥205 SNPs different between human and cattle isolates; **Figure 3** insert); the others were ST540 (929 SNPs different), ST58 (≥1388 SNPs different) and ST69 (≥831 SNPs different). In contrast, there was clear evidence of recent farm-to-farm transmission of isolates from multiple STs (e.g. ST10 and ST69 where, in both cases, there was a 3 SNP minimum distance between pairs of isolates representing two farms) (**Figure 3**). AmpC hyper-producing *E. coli* isolates from these same farms showed a similar pattern: no evidence of strain sharing between dairy farms and humans but strong evidence for recent farm-to-farm transmission (19). We conclude therefore that if humans encounter 3GC-R isolates from farms, such isolates do not readily go on to colonise them and/or cause UTI.

**Figure 3.**
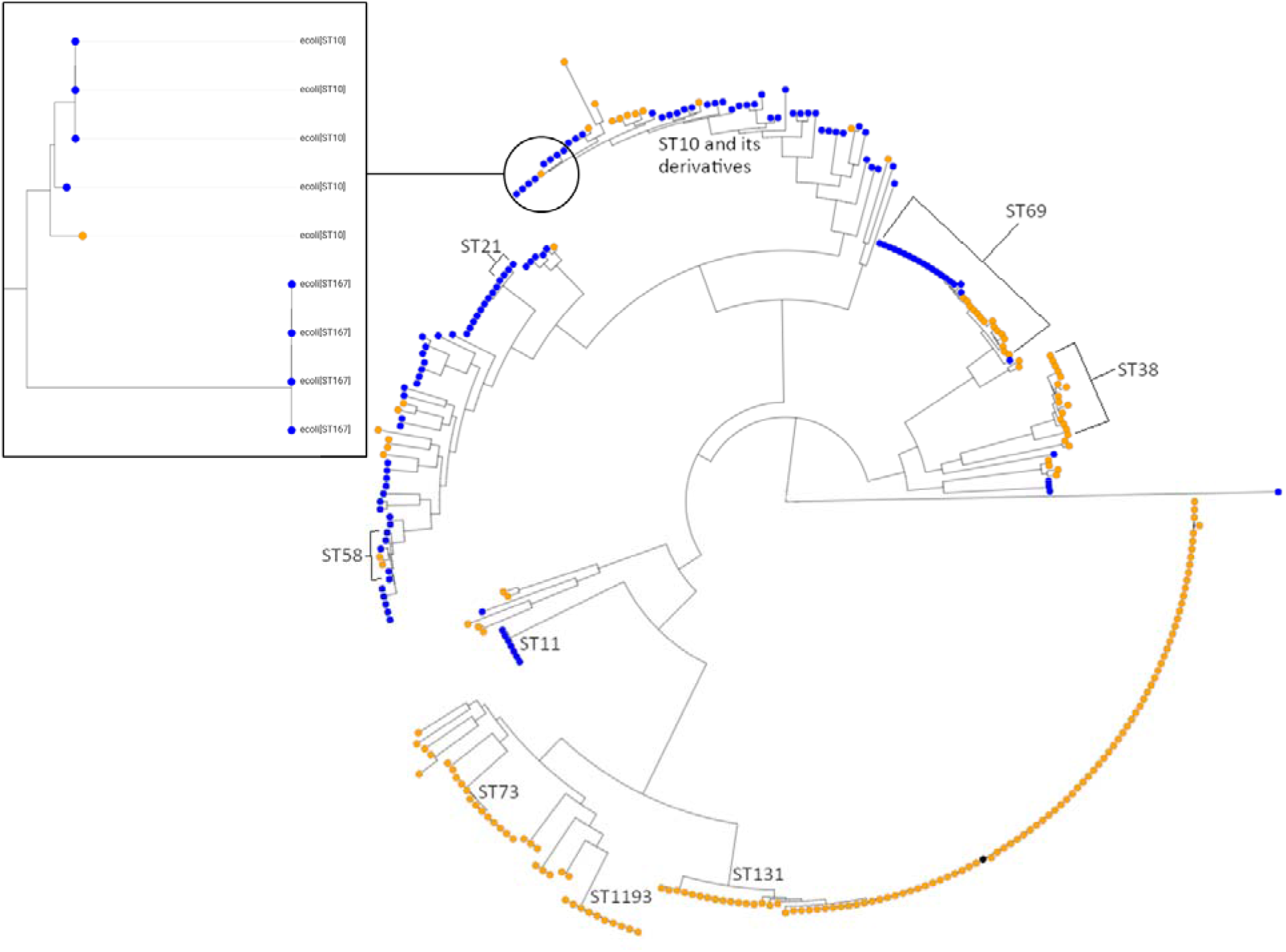
Phylogenetic analysis of *E. coli* from dairy farms and human UTI collected in parallel in a 50 × 50 km region. Human isolates are noted in orange; cattle isolates are noted in blue. The reference ST131 isolate is noted in black. Certain key STs are highlighted, particularly STs with representatives from human and cattle isolates: ST21 (ST540), ST69, ST58 and ST10. The insert shows a more detailed analysis of ST10 isolates which represents the closest relationship between a human and a cattle isolate: 205 SNPs different across the core genome.

We next considered transmission of plasmid-mediated 3GC-R between farm and human *E. coli* isolates (i.e. those that that have caused UTI). We found that 37/107 *bla*_CTX-M_-positive *E. coli* isolates from farm samples within our 50 × 50 km study region carried *bla*_CTX-M_ variants also seen amongst 189 *bla*_CTX-M_-positive urinary *E. coli* cultured from people living in the same region during the same time period (20). None of the human isolates carried *bla*_CTX-M-32_ or any plasmid related to pMOO-32. By filtering sequenced isolates by their *bla*_CTX-M_ variants and plasmid replicon types, plasmids that shared high degrees of sequence identity in farm and human isolates were identified. One IncI1-ST3 plasmid, found in an ST345 farm isolate, harboured *bla*_CTX-M-1_. Mapping of sequencing reads showed that this plasmid exhibited 100% coverage and 97.5% sequence identity to an unpublished ~106 kb IncI1-ST3 plasmid (pTC_N40607; GenBank Accession No. CP007651) found in *E. coli* obtained from meat/cattle isolates in the USA. Six human urinary *E. coli* isolates representing STs 23, 127, 131, 141 and 2015 harboured *bla*_CTX-M-1_ on a plasmid that exhibited 99.4-100% coverage and 96.4-98.7% identity when sequencing reads were mapped to pTC_N40607.

Another plasmid type - again obtained from a single farm isolate, in this case of ST58 - exhibited 100% coverage and 98.5% identity by read mapping to a published IncK plasmid, pCT (GenBank Accession No. NC_014477). pCT is ~94 kb and harbours *bla*_CTX-M-14_. pCT-like plasmids have been reported in both human and veterinary *E. coli* isolates across three continents (12, 23). Amongst human urinary *E. coli* isolates found in this study, two also carried pCT-like plasmids. Both isolates were the pandemic clone ST131, and their pCT-like plasmids exhibited 96.4 or 97.2% identity and 100% coverage to pCT.

In terms of plasmids encoding pAmpC genes, we identified a *bla*_CMY-2_ encoding plasmid, p96 (GenBank Accession no. CP023370). This is a ~96 kb IncI1 plasmid derived from an *E. coli* that caused urinary tract infections in dogs in Scotland (24). Mapping analyses showed that one out of seven *bla*_CMY-2_-carrying human urinary isolates (ST 80) showed 100% coverage and 99.9% identity to p96. An almost identical plasmid was found in one farm isolate (ST1480) that showed 99.6% coverage and 98.9% identity to p96. As with the two *bla*_CTX-M_ plasmids, this *bla*_CMY-2_ encoding plasmid appears to be globally widespread. Blast analysis found *bla*_CMY-2_ plasmids with >95% coverage and >98% identity to p96 in *E. coli* isolates from humans in Taiwan (25) and chicken meat in the USA (GenBank Accession no. CP048295).

### Concluding remarks

Overall, this analysis finds no evidence of recent, direct sharing of *E. coli* between farms and the local human population that have resulted in UTI. However, three farm *E. coli* isolates carried a *bla*_CTX-M_ or *bla*_CMY-2_ plasmid almost identical to one of three plasmids found in urinary *E. coli* in the local human population. However, these three plasmids are known to be widely disseminated in humans and animals across several continents. Furthermore, no human/farm plasmid pair shared 100% identity. So, whilst there is some general evidence of shared circulating plasmids, as reported in a number of recent studies (14,26,27), the overall level of overlap between UK dairy farm and human 3GC-R *E. coli* identified in this study was very small and not suggestive of any novel or recent zoonotic transmission events. In contrast, both recent and sustained farm-to-farm transmission of 3GC-R *E. coli*, and particularly of a newly identified epidemic plasmid pMOO-32 across many different *E. coli* STs and dairy farms, was clearly seen. Identifying the vectors for this transmission will inform interventions that might facilitate a more rapid reduction in 3GC-R *E. coli* on dairy farms.

## Experimental

### Bacterial isolates, identification, and susceptibility testing

Details of farm sample collection and microbiological analysis has recently been reported (18). In brief, samples of faecally contaminated sites were collected using sterile overshoes on 53 dairy farms located in South West England between January 2017 and December 2018. Samples were plated onto TBX agar (Sigma-Aldrich, Poole, UK) containing 16 mg.L^−1^ cephalexin. Up to 5 *E. coli* colonies per cefalexin plate were re-plated onto TBX agar containing 2 mg.L^−1^ cefotaxime to confirm resistance. Presence of acquired 3GC-R genes (*bla*_CTX-M_, *bla*_SHV_, *bla*_CMY-2_ and *bla*_DHA-_ 1) was confirmed by PCR. Disc susceptibility testing was performed and interpreted according to EUCAST guidelines (28).

### Transconjugations

Transconjugations were performed using rifampicin-resistant (Rif-R) *E. coli* DH5α with both human and cattle *E. coli* isolates as the recipients (**Table 3**). Briefly, 1 mL each of overnight broth cultures of donor and recipient cells were mixed in a 3:1 ratio before centrifugation and resuspension in 50 μL of PBS. Five microlitre aliquots were spotted onto LB agar (Oxoid, Basingstoke, UK) plates and incubated at 37°C for 6 h. Growth was collected and resuspended in 100 μL of PBS before being plated on MacConkey agar (Oxoid) plates containing either 32 mg.L^−1^ rifampicin (for Rif-R *E. coli* DH5α) or 0.5 mg.L^−1^ ciprofloxacin (for strains HC4 and HG), and 2 mg.L^−1^ cefotaxime. Transconjugant colonies were screened by PCR.

**Table 3.** Characteristics of *E. coli* strains used in transconjugation experiments.

| Isolate | Source/Host | ST | Resistance Genes | Use |
| --- | --- | --- | --- | --- |
| DK | Cattle | 155 | *strA*, *strB*, *aph(6)-Ic*, *aph(3')-IIa*, *tet(B)*, *bla*<sub>CTX-M-32</sub> | Donor |
| HC4 | Human | 1193 | *aadA5*, *dfrA17*, *mdf(A)*, *sul1* | Recipient |
| HG | Cattle | 88 | *aph(6)-Id*, *ant(2'')-Ia*, *aph(3')-Ia*, *aadA24*, *aph(3'')-Ib*,<br>*sul1*, *sul2*, *tet(A)*, *tet(B)*, *bla*<sub>TEM-1</sub>, *bla*<sub>OXA-1</sub>, *catA1*,<br>*floR* | Recipient |

### WGS and analyses and pMOO-32 PCR

Representative isolates were selected for WGS based on resistance phenotype, β-lactamase gene carriage and farm of isolation, as defined previously (18). WGS was performed by MicrobesNG (https://microbesng.uk/) on a HiSeq 2500 instrument (Illumina, San Diego, CA, USA) using 2×250 bp paired end reads. Reads were trimmed using Trimmomatic (29) and assembled into contigs using SPAdes 3.13.0 (30) (http://cab.spbu.ru/software/spades/). Resistance genes, plasmid replicon types and sequence types (according to the Achtman scheme [31]) were assigned using the ResFinder (32), PlasmidFinder (33), and MLST 2.0 on the Center for Genomic Epidemiology (http://www.genomicepidemiology.org/) platform. Enhanced genome sequencing (combining Illumina and MinION reads) was performed by MicrobesNG on one transconjugant and reads were assembled using Unicycler (34). Contigs were annotated using Prokka 1.2 (35).

Reads and assembled contigs were aligned to reference sequences using the progressive Mauve alignment software (36), CLC Genomics Workbench 12 (Qiagen, Manchester, UK), or BWA (37) and SAMtools (38), with variant positions being called using BCFtools (39). pMOO-32 was visualised using the CGView server (40) (http://stothard.afns.ualberta.ca/cgview_server/).

A multiplex PCR, targeting five size-distinguishable regions of pMOO-32, was designed to indicate the presence of pMOO-32-like plasmids (**Table 4**).

**Table 4.**
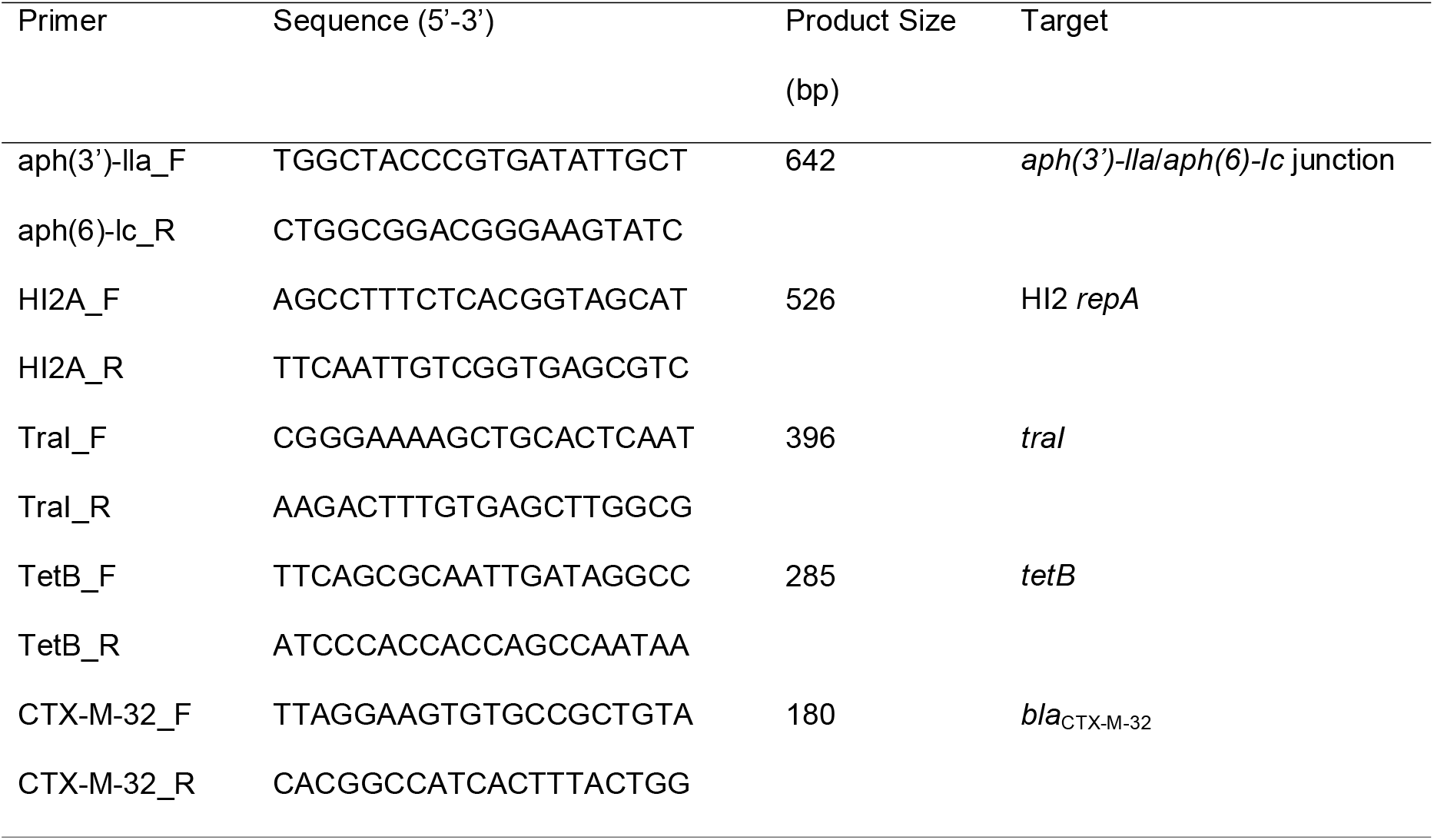
Primers used for the pMOO-32 multiplex PCR.

### Plasmid stability assay

Three representative pMOO-32 PCR-positive isolates, obtained from different farms, and their transconjugant counterparts were subjected to 10 days of serial passaging on non-selective LB agar. After 10 days, colonies were screened for the presence of pMOO-32 by PCR.

### Fitness cost assay

Fitness cost was assessed by a growth curve assay using M9 minimal medium (Sigma-Aldrich). Rif-R *E. coli* DH5α and the pMOO-32 transconjugant strain were grown with shaking at 37°C and OD_600_ measurements were taken at hourly intervals. Assays were performed on three biological replicates.

### Phylogenetic analysis

Sequence alignment and phylogenetic analysis was carried out using the Bioconda environment (41) on the Cloud Infrastructure for Microbial Bioinformatics (CLIMB) (42). The reference sequence was *E. coli* ST131 isolate EC958 complete genome (accession: HG941718). Sequences were first aligned to a closed reference sequence and analysed for SNP differences, whilst omitting insertion and deletion elements, using the Snippy alignment program. Alignment was then focused on regions of the genome found across all isolates, the core genome, using the Snippy-core program, thus eliminating the complicating factors of insertions and deletions (https://github.com/tseemann/snippy). Aligned sequences were then used to construct a maximum likelihood phylogenetic tree using RAxML, utilising the GTRCAT model of rate heterogeneity and the software’s autoMR and rapid bootstrap to find the best-scoring maximum likelihood tree and including tree branch lengths, defined as the number of base substitutions per site compared (43,44). Finally, phylogenetic trees were illustrated using the web-based Microreact program (45).

## Acknowledgements

Genome sequencing was provided by MicrobesNG (http://www.microbesng.uk). We wish to thank all the farmers who participated in this study.

## Transparency declaration

The authors declare no conflict of interests. Farming businesses who permitted access to collect the isolates studied here were not involved in the design of this study or in data analysis and were not involved in drafting the manuscript for publication.

## Author Contributions

Conceived the Study: K.K.R., M.B.A.

Collection of Data: J.F., N.N., O.M., K.M., supervised by T.A.C., M.B.A.

Cleaning and Analysis of Data: J.F., O.M., W.L., H.S., V.C.G., supervised by K.K.R., M.B.A.

Initial Drafting of Manuscript: J.F., M.B.A. Corrected and Approved Manuscript: All Authors

